# Turning coldspots into hotspots: targeted recruitment of axis protein Hop1 stimulates meiotic recombination in *Saccharomyces cerevisiae*

**DOI:** 10.1101/2022.05.12.491616

**Authors:** Anura Shodhan, Martin Xaver, David Wheeler, Michael Lichten

## Abstract

The DNA double strand breaks (DSBs) that initiate meiotic recombination are formed in the context of the meiotic chromosome axis, which in budding yeast contains a meiosis-specific cohesin isoform and the meiosis-specific proteins Hop1 and Red1. Hop1 and Red are important for DSB formation; DSB levels are reduced in their absence and their levels, which vary along the lengths of chromosomes, are positively correlated with DSB levels. How axis protein levels influence DSB formation and recombination remains unclear. To address this question, we developed a novel approach that uses a bacterial ParB-*parS* partition system to recruit axis proteins at high levels to inserts at recombination coldspots where Hop1 and Red1 levels are normally low. Recruiting Hop1 markedly increased DSBs and homologous recombination at target loci, to levels equivalent to those observed at endogenous recombination hotspots. This local increase in DSBs did not require Red1 or the meiosis-specific cohesin component Rec8, indicating that, of the axis proteins, Hop1 is sufficient to promote DSB formation. However, while most crossovers at endogenous recombination hotspots are formed by the meiosis-specific MutLγ resolvase, only a small fraction of crossovers that formed at an insert locus required MutLγ, regardless of whether or not Hop1 was recruited to that locus. Thus, while local Hop1 levels determine local DSB levels, the recombination pathways that repair these breaks can be determined by other factors, raising the intriguing possibility that different recombination pathways operate in different parts of the genome.

## INTRODUCTION

During meiosis, the diploid genome is reduced by half to form haploid gametes by the separation of homologous chromosomes of different parental origin (herein called homologs) during the first of two nuclear divisions (meiosis I). Faithful segregation of homologs requires that they must first identify and link with each other. This is achieved by homologous recombination, which first promotes homolog pairing and then forms crossovers that physically connect homologs and ensure their proper disjunction at meiosis I (Zickler and Kleckner 1999; Whitby 2005; Ur and Corbett 2021). Errors in homologous recombination cause aneuploidy in gametes, which in turn causes infertility, pregnancy loss, and genetic disorders (Hassold and Hunt 2001; Srivastava *et al*. 2012; Wang *et al*. 2017; Gao *et al*. 2018).

Meiotic recombination occurs in the context of a chromosome axis that contains three components: cohesin; an axis core protein; and a HORMA domain-containing protein (Hollingsworth and Ponte 1997; Zickler and Kleckner 1999; Blat *et al*. 2002; Glynn *et al*. 2004; Tsubouchi and Roeder 2006; Niu *et al*. 2007; Yang *et al*. 2008; Kugou *et al*. 2009; Niu *et al*. 2009; Callender and Hollingsworth 2010; Kim *et al*. 2010; Panizza *et al*. 2011; Chuang *et al*. 2012; Pyatnitskaya *et al*. 2019; Ur and Corbett 2021). The cohesin core holds the sister chromatids together and organizes them in a linear array of loops (Smith and Roeder 1997; Zickler and Kleckner 1999; van Heemst and Heyting 2000; Kleckner 2006; Lam and Keeney 2014). Meiotic cohesin, which contains the meiosis-specific kleisin subunit Rec8, is important for most of the chromosomal localization of the other two axis proteins in wild type cells (Smith and Roeder 1997; Klein *et al*. 1999; Blat *et al*. 2002; Riedel *et al*. 2006; Jin *et al*. 2009; Joshi *et al*. 2009; Katis *et al*. 2010; Panizza *et al*. 2011; Sun *et al*. 2015; Heldrich *et al*. 2020; Heldrich *et al*. 2022). Axis core proteins (Red1 in *S. cerevisiae*, ASY3/4 in *Arabidopsis*, SYCP2/3 in mammals, Rec10/27 in *S. pombe*) have diverged considerably in sequence but have similar domain structures and are functionally conserved (Rockmill and Roeder 1990; Hollingsworth and Ponte 1997; Smith and Roeder 1997; de los Santos and Hollingsworth 1999; West *et al*. 2019; Ur and Corbett 2021). HORMA domain-containing proteins (Hop1 in *S. cerevisiae* and *S. pombe*, ASY1/2 in *Arabidopsis*, HORMAD1/2 in mammals, HTP-1/2/3/HIM-3 in *C. elegans*) are highly conserved, and in most organisms contain a HORMA domain and a loop containing a peptide sequence, called a closure motif, that binds either to its own HORMA domain to form a closed structure or to a HORMA domain on another protein to form oligomers (Moses 1956; Hollingsworth and Byers 1989; Hollingsworth and Ponte 1997; Woltering *et al*. 2000; Martinez-Perez and Villeneuve 2005; Yang *et al*. 2006; Baudat and de Massy 2007; West *et al*. 2018). HORMA domain proteins are recruited to the axis by an interaction between their HORMA domain and a closure motif on the axis core protein (West *et al*. 2018; West *et al*. 2019). Although the main function of these proteins is similar in most organisms, there are also differences that have been discussed in detail elsewhere (Zickler and Kleckner 2015; Zickler and Kleckner 2016; Ur and Corbett 2021). For simplicity, the rest of this introduction will focus on the function of these proteins in meiotic recombination in *Saccharomyces cerevisiae*.

Chromosome axis proteins are important for the first step of meiotic recombination, the formation of programmed DNA double strand breaks (DSBs) by the meiosis-specific protein Spo11 and its co-factors: the RMM complex (Rec114, Mer2, Mei4); the MRX complex (Mre11, Rad50, Xrs2); Rec102-Rec104; and Ski8 (Malone *et al*. 1991; Bergerat *et al*. 1997; Uetz *et al*. 2000; Keeney 2001; Kee and Keeney 2002; Tesse *et al*. 2003; Arora *et al*. 2004; Kee *et al*. 2004; Prieler *et al*. 2005; Henderson *et al*. 2006; Li *et al*. 2006; Maleki *et al*. 2007; Panizza *et al*. 2011; Stanzione *et al*. 2016). On a regional scale (on the order of 20-50kb), enrichment levels for Spo11 and DSBs are closely related to those observed for Hop1 and Red1 (Hollingsworth and Ponte 1997; Blat *et al*. 2002; Pan *et al*. 2011; Panizza *et al*. 2011; Smagulova *et al*. 2011; Sun *et al*. 2015). In addition, mutant analyses have shown that the absence of any of the axis proteins results in a reduction in DSBs, although the extent of reduction can differ between genome regions (Zickler and Kleckner 1999; Blat *et al*. 2002; Glynn *et al*. 2004; Kugou *et al*. 2009; Kim *et al*. 2010; Panizza *et al*. 2011; Ur and Corbett 2021). *hop1* mutants seem to show the most pronounced DSB reduction, at least when measured at loci where DSBs form frequently, called hotspots (Mao-Draayer *et al*. 1996; Schwacha and Kleckner 1997; Xu *et al*. 1997; Woltering *et al*. 2000; Pecina *et al*. 2002; Niu *et al*. 2005). Hop1 is thought to promote DSB formation by interacting with Mer2, a member of the trimeric RMM complex, and this interaction is conserved in other species (Stanzione *et al*. 2016; Kariyazono *et al*. 2019; Claeys Bouuaert *et al*. 2021; Rousova *et al*. 2021). Mer2, in turn, interacts with the other RMM components as well as other proteins that are important for Spo11-mediated DSB formation (Acquaviva *et al*. 2013; Sommermeyer *et al*. 2013; Rousova *et al*. 2021). *In vitro* studies indicate that, although Red1 has no detectable affinity for Mer2, Red1 stimulates Hop1-Mer2 interaction by changing Hop1’s conformation and increasing its affinity for Mer2 (Rousova *et al*. 2021). Hop1 is also required for cohesin-independent enrichment of Red1 in certain parts of the genome (Panizza *et al*. 2011; Sun *et al*. 2015; Heldrich *et al*. 2020). Taken together, these observations suggest that Hop1 may be the primary axis protein promoting DSB formation, although this has not been directly demonstrated.

Once DSBs form, Hop1 and Red1 play subsequent roles in promoting interhomolog recombination and in promoting crossover formation. DSBs promote Hop1 phosphorylation by the Mec1(ATR)/Tel1(ATM) kinases (Carballo *et al*. 2008), and this promotes use of the homolog rather than the sister chromatid as a repair template (Hollingsworth and Ponte 1997; Tsubouchi and Roeder 2006; Niu *et al*. 2007; Niu *et al*. 2009; Callender and Hollingsworth 2010; Chuang *et al*. 2012). Once paired, homologs are held together by a tripartite proteinaceous structure called the synaptonemal complex and Hop1 is removed from the chromosome axis, curbing further DSB formation and removing the inter-sister recombination barrier to allow quick repair of any remaining breaks (Borner *et al*. 2008; Joshi *et al*. 2009; Wojtasz *et al*. 2009; Zanders and Alani 2009; Daniel *et al*. 2011; Kauppi *et al*. 2013; Thacker *et al*. 2014; Lambing *et al*. 2015; Subramanian *et al*. 2016; Subramanian *et al*. 2019). Red1 interacts with Zip4 (Yang *et al*. 2008; De Muyt *et al*. 2018; Pyatnitskaya *et al*. 2019), a member of the ZMM protein complex (Zip1, Zip3, the Zip2-Zip4-Spo16 complex, the Msh4-Msh5 complex, and Mer3) that stabilizes double Holliday junction intermediates and directs them toward resolution as crossovers by the meiosis-specific resolvase, MutLγ (Mlh1-Mlh3 and Exo1) (Schwacha and Kleckner 1994; Wang *et al*. 1999; Khazanehdari and Borts 2000; Kirkpatrick *et al*. 2000; Tsubouchi and Ogawa 2000; Allers and Lichten 2001b; Allers and Lichten 2001a; Hoffmann *et al*. 2003; Bishop and Zickler 2004; Borner *et al*. 2004; Jessop *et al*. 2006; Lynn *et al*. 2007; Nishant *et al*. 2008; Zakharyevich *et al*. 2010; Comeron *et al*. 2012; Wang *et al*. 2012; Yang *et al*. 2012; Al-Sweel *et al*. 2017; De Muyt *et al*. 2018; Pyatnitskaya *et al*. 2019; Cannavo *et al*. 2020; Kulkarni *et al*. 2020; Sanchez *et al*. 2020). This is the major pathway for crossover formation; a minority of crossovers are formed by the mitotic structure-selective nucleases (SSNs) Mus81-Mms4, Slx1-Slx4, Yen1 (De los Santos *et al*. 2003; Argueso *et al*. 2004; Hollingsworth and Brill 2004; Lynn *et al*. 2007; Jessop and Lichten 2008; De Muyt *et al*. 2012; Zakharyevich *et al*. 2012; Agostinho *et al*. 2013; Oke *et al*. 2014). Joint molecule resolution and crossover formation in both pathways depend on the meiosis-specific transcription factor Ndt80, which drives the mid-meiosis expression of many proteins required to complete meiosis and sporulation, including the polo-like kinase Cdc5 that stimulates resolvase activities (Xu *et al*. 1995; Chu and Herskowitz 1998; Allers and Lichten 2001a; Clyne *et al*. 2003; Sourirajan and Lichten 2008; De Muyt *et al*. 2012; Sanchez *et al*. 2020).

In summary, meiotic axis proteins play roles in various stages of meiotic recombination, with current data indicating that Hop1 has an early role in DSB formation and partner choice, while Red1 has a later role in recombination pathway choice. However, because Red1 and Hop1 are co-dependent for localization, determining the specific role that each protein plays in meiotic recombination remains a challenge. Here, we used a novel approach based on a bacterial ParB-*parS* partition system (Khare *et al*. 2004; Dubarry *et al*. 2006; Murray *et al*. 2006; Sullivan *et al*. 2009; Graham *et al*. 2014; Saad *et al*. 2014; Attaiech *et al*. 2015), to recruit Hop1 to regions where meiotic axis proteins are normally depleted. We find that recruiting Hop1 at high levels is sufficient to dramatically increase both DSBs and homologous recombination, consistent with Hop1 being the most immediate determinant of where meiotic recombination occurs in the genome.

## MATERIALS AND METHODS

### Yeast strains

All *S. cerevisiae* strains (File S1 sheet 1) used in this study are of SK1 background (Kane and Roth 1974) and were made by transformation or genetic crosses. To monitor the effect of axis protein recruitment via the ParB-*parS* system, two recombination reporter inserts were used (for schematics, see Figure 2A and Figure 9A, below). The first is a modification of the previously described *URA3-ARG4-pBR322* insert (Wu and Lichten 1995; Borde *et al*. 1999), and the second a modification of the previously described *URA3-tel-ARG4* insert (Jessop *et al*. 2005; Ahuja *et al*. 2021). For both inserts, a 1kb fragment containing the *parSc2* element from chromosome *c2* of *Burkholderia cenocepacia* J231 (Saad *et al*. 2014) was synthesized and added downstream of the *ARG4* gene. The *URA3-ARG4-pBR322-parS* construct was linearized by *EcoRI* and inserted 237nt downstream of *HXT1* and 150nt downstream of *YCR017c* by ends-out three-piece transformation (primers in File S1 sheet 2). For insertion at *URA3*, the construct was linearized by *ApaI*, which cuts in the *URA3* gene, and was inserted via ends-in one-piece transformation. Hop1-ParB fusions are illustrated in Figure 1A. Sequences encoding ParBc2, which binds to *parSc2* (Saad *et al*. 2014), were modified to include a V5 tag (Funakoshi and Hochstrasser 2009) and a stop codon at its C-terminus. This was combined with *HOP1* flanking sequences in the following order to make pMJ1088 (sequence in File S3): the *HOP1* promoter (+652 to -1 nt); ParBc2-V5—stop codon; *HOP1* 3’UTR (131bp starting at the 3’ end of *HOP1* coding sequences); *natMX4* (Lorenz 2015). PCR products (primers in File S1 sheet 2) containing this element were integrated at *HOP1* by single ends-in transformation to produce a *HOP1* duplication where one copy was C-terminally tagged *[HOP1-parBsc2-V5]-natMX-HOP1]*, and by ends-out replacement transformation to produce a single C-terminally tagged copy of *HOP1* (*[HOP1-parBsc2-V5]-natMX*). Although both *HOP1-V5-parBsc2* and *HOP1* are expressed from the endogenous *HOP1* promoter, levels of the Hop1-V5-ParB fusion protein were about 10-20% lower than of the corresponding wild-type Hop1 protein (Figure 1B, File S1 sheet 14).

**Figure 1.**
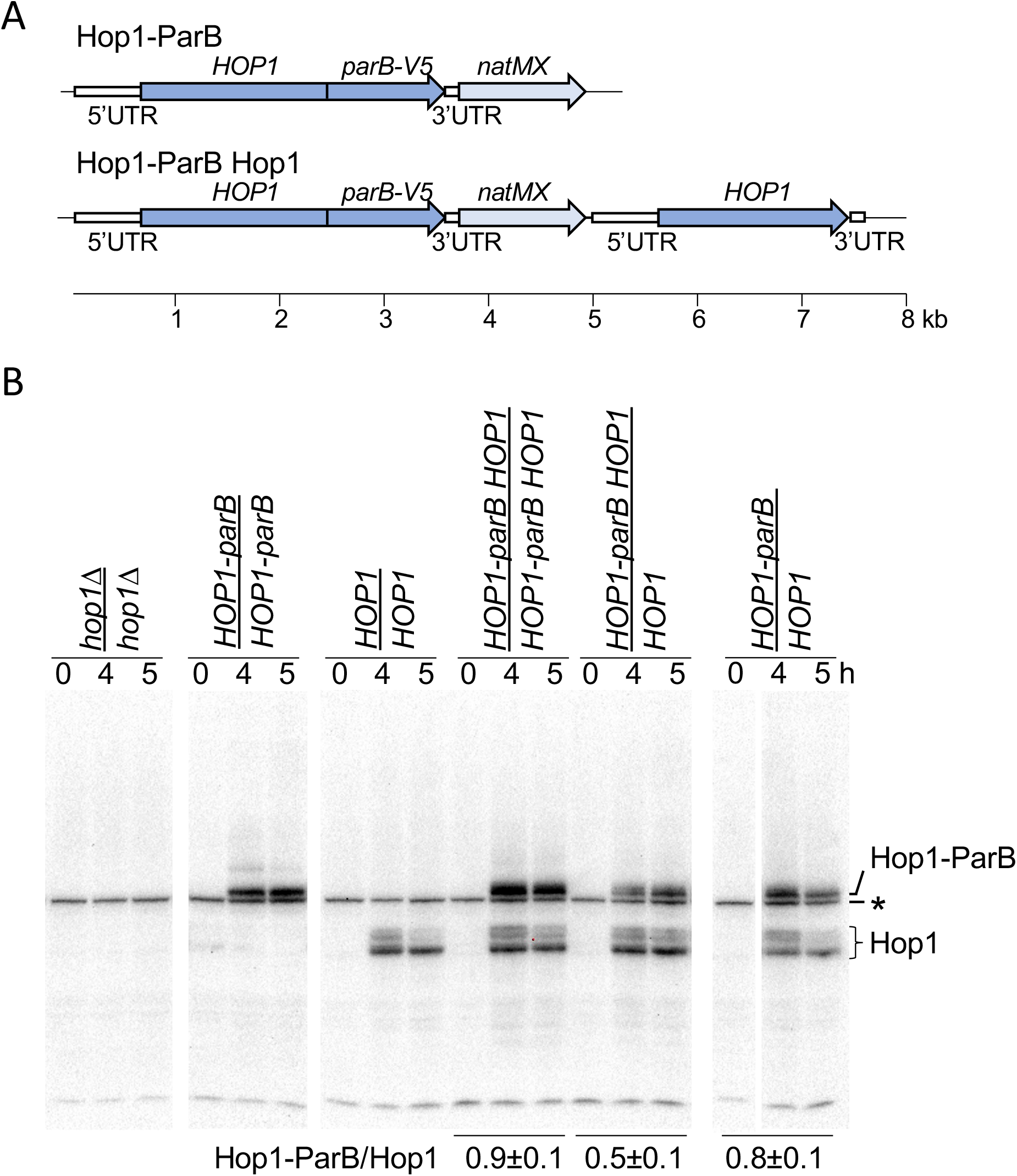
ParB fusion constructs. A. Illustration of protein fusions used. Dark blue arrows –coding sequences of tagged and untagged *HOP1*; vertical lines—fusion; light blue—*natMX6* drug resistance cassette; open boxes—5’ and 3’ *HOP1* untranslated regions; thin rectangles—pFA6 sequences; thin black lines—flanking yeast chromosome sequences. B. Western blot of samples taken at indicated time in meiosis, probed with anti-Hop1. Bands corresponding to Hop1 and to Hop1-ParB are indicated; asterisk indicates non-specific background band. Ratios of Hop1-ParB/Hop1 are indicated for strains where the two proteins are both present. See also File S1 sheet 14.

**Figure 2.**
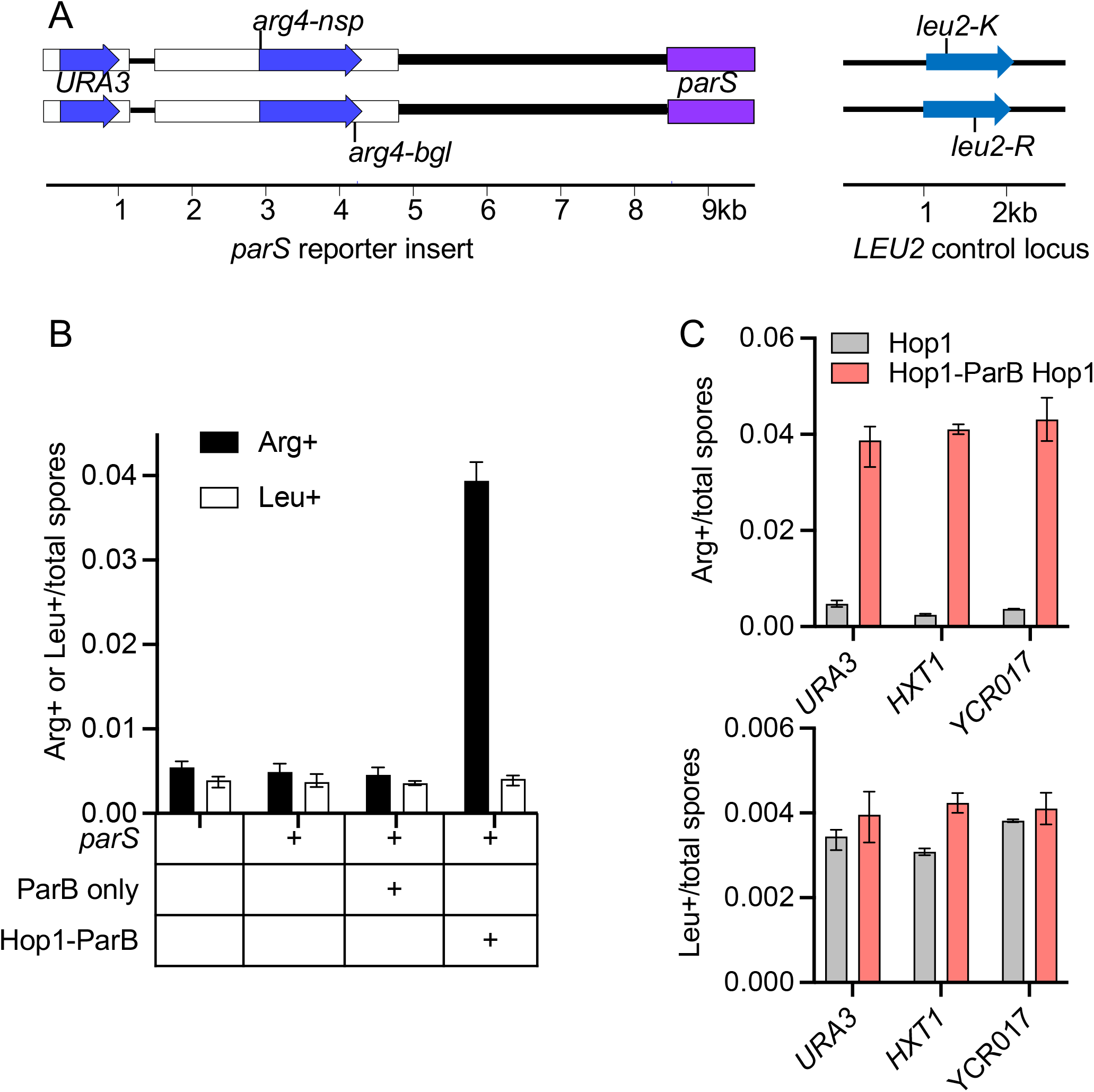
Hop1 recruitment stimulates meiotic recombination. A. Left—schematic of the *URA3-arg4-parS* reporter insert, showing *arg4-nsp* and *arg4-bgl* heteroalleles; right—*leu2* control locus with heteroalleles. Blue arrows—coding sequences; white boxes—yeast chromosomal sequences; purple box—*parSc2* sequences; thick line—pBR322 sequences. B. Frequencies of Arg+ (insert, black) and Leu+ (control, white) recombinants for the insert at *URA3* are shown in A. Fusion proteins expressed are indicated; all strains also expressed wild-type Hop1. C. Frequencies of Arg+ (top) and Leu+ (bottom) recombinants in strains with inserts at the indicated locus, expressing only Hop1 (grey) or both Hop1-ParB and Hop1 (salmon). See also File S1 sheet 4.

To genetically monitor crossovers, markers flanking the *URA3-arg4-pBR322-parS* insert at *URA3* were inserted by transformation (primers in File S1 sheet 2): *kanMX6* (Lorenz 2015) into the intergenic region between *RIP1* and *YEL023c* ∼10kb to the right of the insert; and *hygMX6* (Saad *et al*. 2014) into the intergenic region between *NPP2* and *EDC3* ∼10kb to the left.

### Sporulation, DNA extraction and Southern blots

Strains were grown in liquid pre-sporulation medium and transferred to liquid sporulation medium as described (Goyon and Lichten 1993). Culture samples were collected and processed as described (Allers and Lichten 2000; Jessop *et al*. 2005; Jessop *et al*. 2006). DNA was extracted as described (Goyon and Lichten 1993), digested with the appropriate restriction enzymes, displayed on agarose gels, transferred to membranes, hybridized to radioactive probes (File S1 sheet 3) and analyzed as described (Wu and Lichten 1994; Wu and Lichten 1995; Allers and Lichten 2001b).

### Western blots

Protein was extracted from meiotic cultures, displayed on polyacrylamide gels, blotted to membranes and probed basically as described (Kaur *et al*. 2018), except that nonfat dry milk was used as in place of iBlock. Primary antisera and dilutions used were: rabbit anti-Hop1 (made for this work, 1:75000) and goat anti-Arp7 ((Santa Cruz Biotechnology Cat# sc-8961, RRID:AB_671730, 1:1000). Secondary antisera were: goat polyclonal anti-rabbit conjugated with alkaline phosphatase (Abcam Cat# ab97048, RRID:AB_10680574, 1:10000) and rabbit anti-goat IgG conjugated with alkaline phosphatase (Sigma-Aldrich Cat# A4187, RRID:AB_258141, 1:5000). Chemiluminescence signals were captured using a BioRad Chemidoc MP imaging system and were quantified using the gel quantification tools in Fiji (Schindelin *et al*. 2012).

### Cytology

Nuclear divisions were monitored by DAPI staining as described (Goyon and Lichten 1993). Meiotic chromosome spreads and staining with antisera were performed as described (Loidl *et al*. 1991). The primary antibodies were: rabbit polyclonal anti-Hop1 serum (prepared for this project), 1:7500 and mouse anti-V5 (Bio-Rad Cat# MCA1360, RRID:AB_322378, 1:250). The secondary antibodies were: goat anti-rabbit conjugated to Alexa 488 (Molecular Probes, #A-11034), 1:350 and donkey anti-mouse conjugated to Cy3 (Jackson ImmunoResearch Labs Cat# 715-165-151, RRID:AB_2315777, 1:500). Images were taken on a Zeiss Axioplan 2 imaging microscope using a 100x plan apochromat objective (440782-9902) and a Zeiss AxioCam HRm camera.

### Genetic analysis

Frequencies of recombination between heteroalleles were determined by random spore analysis as described (Lichten *et al*. 1987). Map distances were determined by tetrad dissection, using the formula of Perkins (Perkins 1949) as implemented at https://elizabethhousworth.com/StahlLabOnlineTools/compare2.php.

### Calibrated chromatin immunoprecipitation and sequencing (ChIP-seq)

ChIP-seq experiments used a protocol that combined and modified previous methods ((Murakami and Keeney 2014; Makrantoni *et al*. 2019); Hajime Murakami, personal communication). Strains used contained the *URA3-tel-ARG4-parS* reporter construct inserted at *URA3*. Samples taken at 0, 3 and 4h post meiotic induction were fixed with 1% formaldehyde for 30 min at room temperature and quenched with 125mM glycine. The cells were washed in 1X TBS (20 mM Tris-HCl, pH 7.5, 136 mM NaCl) and stored as a pellet at -80°C. *Saccharomyces mikatae* cells were similarly fixed 4h post meiotic induction and aliquots were frozen that contained about 1/10^th^the number of cells taken for *S. cerevisiae*. Both pellets were mixed in 500µl lysis buffer (50 mM Hepes-KOH pH 7.5, 140 mM NaCl, 1 mM EDTA, 1% Triton X-100, 0.1% sodium deoxycholate, 1X Complete Protease Inhibitor Cocktail EDTA-free (Roche, #04693132001), 7µg/ml aprotinin (Thermo Scientific, #78432), 1mM PMSF) and lysed in a Mini-Beadbeater-16 (Biospec products) for seven cycles of 1min on, 2 min off (where the samples were kept on ice). The lysate was sonicated using a Biorupter 300 (Diagnode) for two rounds of eleven cycles of 30 sec on/30 sec off, with a 20 min incubation on ice between the two rounds of sonication. Debris was then removed by centrifugation (21130 x g, 5min, 4°C) and another round of 11 cycles of sonication was performed. Lysates were pre-cleared by incubating with 50µl protein G-conjugated Dynabeads (Invitrogen, #100.04D; beads were washed twice with 1ml lysis buffer before use) for 1h on a rotator at 4°C. Beads were removed, and a 10µl sample of the lysate was mixed with 190µl 10mM TRIS, 1mM EDTA, 1% SDS pH7.5 and stored at 4°C to be used as input DNA. 3µl of anti-Hop1 serum was added to the remaining lysate, which was then incubated for 3h at 4°C with rotation. Protein G-conjugated Dynabeads (50µl, washed twice with 400µl 137 mM NaCl, 2.7 mM KCl, 10 mM Na_2_HPO_4_, 1.76 mM KH_2_PO_4_, pH 7.4 with 5mg/ml bovine bovine serum albumin) were added and the mixture was incubated overnight at 4°C with rotation. Beads were then washed twice with 1ml of the following three buffers in succession; lysis buffer, wash buffer I (10 mM Tris-HCl pH 8, 250 mM LiCl, 360mM NaCl, 0.5% Na-deoxycholate, 1 mM EDTA, 0.5% Triton X-100), wash buffer II (10 mM Tris-HCl pH 8, 250 mM LiCl, 0.5% Na-deoxycholate, 1 mM EDTA, 0.5% Triton X-100); for 5 min each on a rotator at 4°C. The beads were washed once with 1ml TE wash buffer (10 mM Tris-HCl pH 8, 1 mM EDTA, 0.5% Triton X-100) at 4°C for 5 min with rotation. DNA was eluted in 40µl elution buffer (50 mM Tris-HCl pH 8, 10 mM EDTA, 1% SDS) at 65°C for 15 min and added to a tube containing 160µl of TE/1%SDS. 200µl of ChIP and input DNA were incubated overnight at 65°C in the presence of 1µl RNAse (0.5 mg/ml) to reverse crosslinks. 7.5µl of proteinase K (20mg/ml) was added to each tube and incubated at 50°C for 2h. DNA was purified using a QIAquick PCR purification kit (Qiagen, #28104) and eluted in 50µl water. 15 ng of ChIP and input DNA were used to generate libraries using NEBNext Ultra II DNA Library Prep Kit for Illumina (New England Biolabs, #E7645) and NEBNext® Multiplex Oligos for Illumina® (96 Unique Dual Index Primer Pairs, New England Biolabs, #E6440). Sequencing was performed with an Illumina NextSeq 550 with the NextSeq 500/550 High Output Kit v2.5 (75 Cycles).

ChIP-seq data were calibrated as described (Makrantoni *et al*. 2019). Briefly, single ended fastq format sequences derived from ChIP-seq data were quality-trimmed using fastp (Chen *et al*. 2018). Trimmed fastqs from both IP and input were aligned separately to the *SK1* target genome ((Yue *et al*. 2017), available at https://yjx1217.github.io/Yeast_PacBio_2016/data/) which had been modified to reflect the genotype of the diploid MJL4236/7 (File S1 sheet1) and also to the *Saccharomyces mikatae* IFO 1815 (Kellis *et al*. 2003) spike-in control genome using minimap2 (Li 2018). Reads that did not map to SK1 were subsequently aligned to *S. mikatae* and vice versa to identify those reads that mapped to both genomes and those that mapped uniquely to a single genome. A calibration factor, called the occupancy ratio (OR), was then calculated from the counts of such reads as:

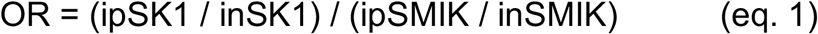

Where,

ipSK1 = count IP reads mapping uniquely to the SK1 genome

inSK1 = count of input reads mapping uniquely to the SK1 genome

ipSMIK = count of IP reads mapping uniquely to the *S. mikatae* genome

inSMIK = count of input reads mapping uniquely to the *S. mikatae* genome

Calibrated depths for reads mapping uniquely to the SK1 genome were determined by multiplying read depths per million mapped reads by the OR computed in equation 1. Data processing was performed on the NIH HPC Biowulf cluster (http://hpc.nih.gov). Scripts implementing the calibrated ChIP processing pipeline as a Snakemake (Molder *et al*. 2021) workflow, suitable for parallel execution on the Biowulf cluster, are included in File S2; sequence reads are available at GEO, accession GSE201240, access token provided upon request.

### Data representation

All values reported in figures are the mean of two or more independent experiments. Error bars denote the range in data values, except in Figure 8, where they denote calculated standard error.

## RESULTS

To recruit axis proteins to target loci, we used the bacterial ParB-*parS* chromosome segregation system, where the ParB protein binds to a <1kb-long cluster of *parS* sites and then spreads to adjacent DNA (Lin and Grossman 1998; Dubarry *et al*. 2006; Breier and Grossman 2007; Attaiech *et al*. 2015; Soh *et al*. 2019). This system allows recruitment of multiple copies of ParB, fused to a protein of interest, with minimal disruption of chromosome integrity and function (Dubarry *et al*. 2006; Saad *et al*. 2014). We fused ParB and a V5 epitope tag to the C-terminus of Hop1 (hereafter called Hop1-ParB; Figure 1A) to target this protein to three loci: *URA3*; *HXT1*; and *YCR017c*. All three are in regions of the yeast genome with low levels of occupancy by meiotic axis proteins and low levels of meiotic DSBs (Figure S1; (Pan *et al*. 2011; Panizza *et al*. 2011)).

### Recruiting Hop1 increases meiotic recombination

To determine the effect of recruiting Hop1 on meiotic recombination, we initially used random spore analysis to examine recombination between *arg4* heteroalleles in a *URA3-arg4-pBR322-parS* recombination reporter inserted at *URA3* (Figure 2A). The same insert, but without *parS*, forms DSBs and undergoes recombination at levels that are location-dependent and that reflect underlying recombination levels in the region where it is inserted (Borde *et al*. 1999). As a non-insert control, we also measured recombination between heteroalleles at *LEU2*. Initial experiments used a *HOP1* geneduplication that contained both a tagged and a wild-type copy, to ensure normal function in the event that the tagged protein was only partially functional.

Recruiting Hop1-ParB caused a striking 7-fold increase in recombination in the *arg4* gene inserted at *URA3* (Figure 2B; File S1 sheet 4). Inserts at *HXT1* and *YCR017c*, two other axis protein/DSB coldspots (Figure S1), also displayed markedly increased Arg+ recombinant frequencies (10-and 15-fold, respectively) when Hop1-ParB was present (Figure 2C). The presence or absence of a ParB-tagged axis protein did not markedly change recombination frequencies at the *leu2* control locus (3.6 ± 0.7 × 10^−3^, Figure 2B,C; File S1 sheet 4). These results suggest that levels of Hop1 in a region might be sufficient to determine levels of meiotic recombination in that region.

Recruiting Hop1-ParB also markedly increased crossing-over in a region containing the insert at *URA3*. Crossing-over was measured by analysis of tetrads from a diploid that contained a *kanMX6* insert ∼10kb centromere proximal to *URA3-arg4-pBR322-parS* in one parent, and a *hygMX6* insert ∼10kb centromere distal in the other parent. The genetic distance for this ∼25 kb interval was 12.3 ± 1.78 cM in diploids lacking Hop1-ParB and 66.2 ± 6.16 cM in diploids expressing Hop1-ParB, a ∼5-fold stimulation of crossing-over (File S1 sheet 10, also see Figure 8, below).

### Recruiting Hop1 increases DSB formation

To confirm that recruiting Hop1 increases meiotic recombination by increasing levels of DSBs, we determined cumulative DSB levels in *sae2Δ* mutants, which accumulate unrepaired DSBs with unresected ends (Keeney and Kleckner 1995; Prinz *et al*. 1997). Consistent with previous data (Borde *et al*. 1999; Pan *et al*. 2011), very few DSBs were present in reporter inserts at the three target loci (*URA3, HXT1* and *YCR017c*) in the absence of Hop1-ParB or when ParB alone was expressed. The presence of Hop1-ParB increased DSBs in the reporter construct dramatically at all three loci (Figures 3A and 3B; File S1 sheet 5), while the DSBs at the *ARE1* control locus (Goldway et al. 1993) were relatively unchanged (Figure 3B). Hop1-ParB recruitment caused the greatest increase in DSBs in the insert at *URA3* locus, where DSB levels (∼21% of chromatids) are consistent with most cells experiencing a break at this locus.

**Figure 3:**
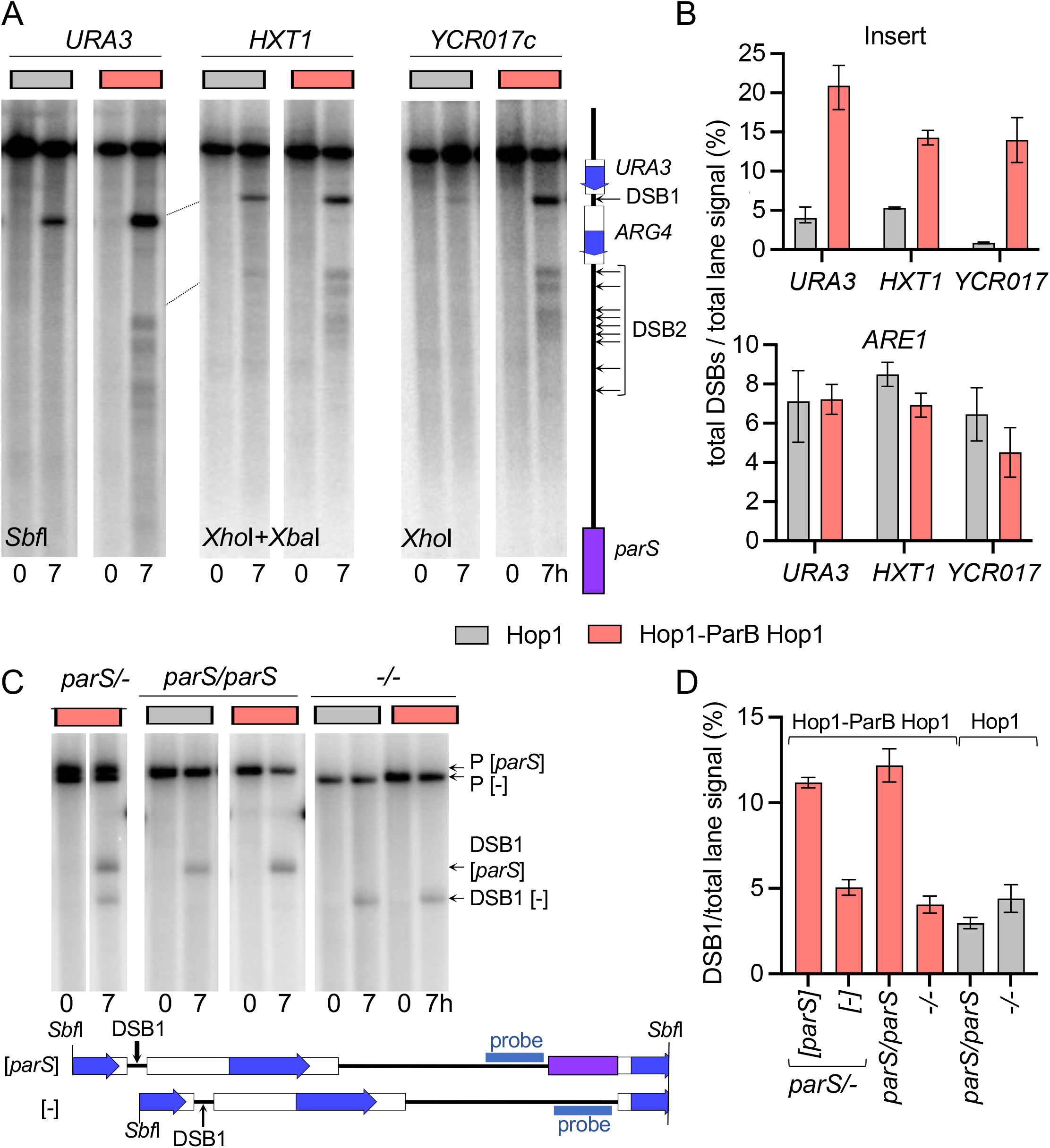
Hop1-ParB recruitment increases DSBs in reporter inserts. A. Southern blot of DNA from *sae2Δ* strains, which form DSBs but do not resect or repair them, with a *parS* insert at indicated locus. Indicated restriction digests were probed with *parS* sequences to detect DSBs in the insert. These occur in pBR322 sequences on either side of *ARG4* sequences (Wu and Lichten 1995), and will be called DSB1 and DSB2 as shown in the schematic. Strains were homozygous either for *HOP1* (grey) or *HOP1-parB HOP1* (salmon). B. Hop1-ParB increases DSBs (DSB1 + DSB2) at all three insert loci (top), but not at the *ARE1* control locus (Bottom). C. Hop1-parB acts primarily in *cis:* Southern blot with DNA from a *sae2Δ* strain with inserts at *URA3* on both homologs, where: *(parS/-*)—one contains *parS* and the other does not; (*parS/parS*)—both contain *parS*; (-/-)—both are without *parS*. DNA was digested with *Sbf*I and probed with pBR322 sequences, which allows distinction between breaks at DSB1 on chromosomes with and without *parS*. Breaks at DSB2 cannot be resolved. D. Quantification of breaks at DSB1 in the *parS* hemizygous strain, as well as in control strains with homozygous inserts that either both lacked or both contained *parS*. See also File S1 sheets 5 and 6.

We also asked if Hop1 levels affect DSBs in *cis* or *trans*. In strains with *parS* on only one of the two homologs, the homolog with *parS* displayed insert DSBs at levels similar to those seen in a *parS*-homozygous diploid, while the homolog lacking *parS* displayed DSBs at levels similar to those seen in strains without *parS* (Figure 3C,D; File S1 sheet 6). Thus, the DSB increase observed is primarily due to recruited Hop1 acting in *cis*.

### Hop1-ParB-stimulated DSBs require Spo11 but not Rec8 or Red1

According to current models, DSBs are formed by the Spo11 complex, which is recruited to the cohesin-based axis by interactions with Hop1, which in turn can be recruited to the axis via interactions with Red1 (Panizza *et al*. 2011; Sun *et al*. 2015; Zickler and Kleckner 2015; West *et al*. 2019; Rousova *et al*. 2021). This suggests that artificially recruiting Hop1 to chromosomes might bypass the need for Red1 or cohesin in DSB formation. To test this suggestion, DSBs in the insert at *URA3* were examined in *sae2Δ* strains that were lacking Spo11, Red1, or the meiosis-specific cohesin component Rec8.

As expected, DSBs were abolished at all loci in *spo11Δ* strains, regardless of whether Hop1-ParB was present (Figures 4A and 4B; File S1 sheet 5). Consistent with previous reports (Woltering *et al*. 2000; Pecina *et al*. 2002; Niu *et al*. 2005), *red1Δ* mutants displayed a substantial (∼2.5-fold) decrease in DSBs at the *ARE1* control locus regardless of whether Hop1-ParB was present or absent; when only Hop1 was present, the *parS* insert locus showed a similar decrease in DSBs. However, when Hop1-ParB was present, DSB levels at the *parS* insert in *red1Δ* strains were similar to those in *RED1* strains (Figures 4A and 4B; File S1 sheet 5). Thus, direct recruitment of Hop1 appears to bypass the role of Red1 in DSB formation.

**Figure 4:**
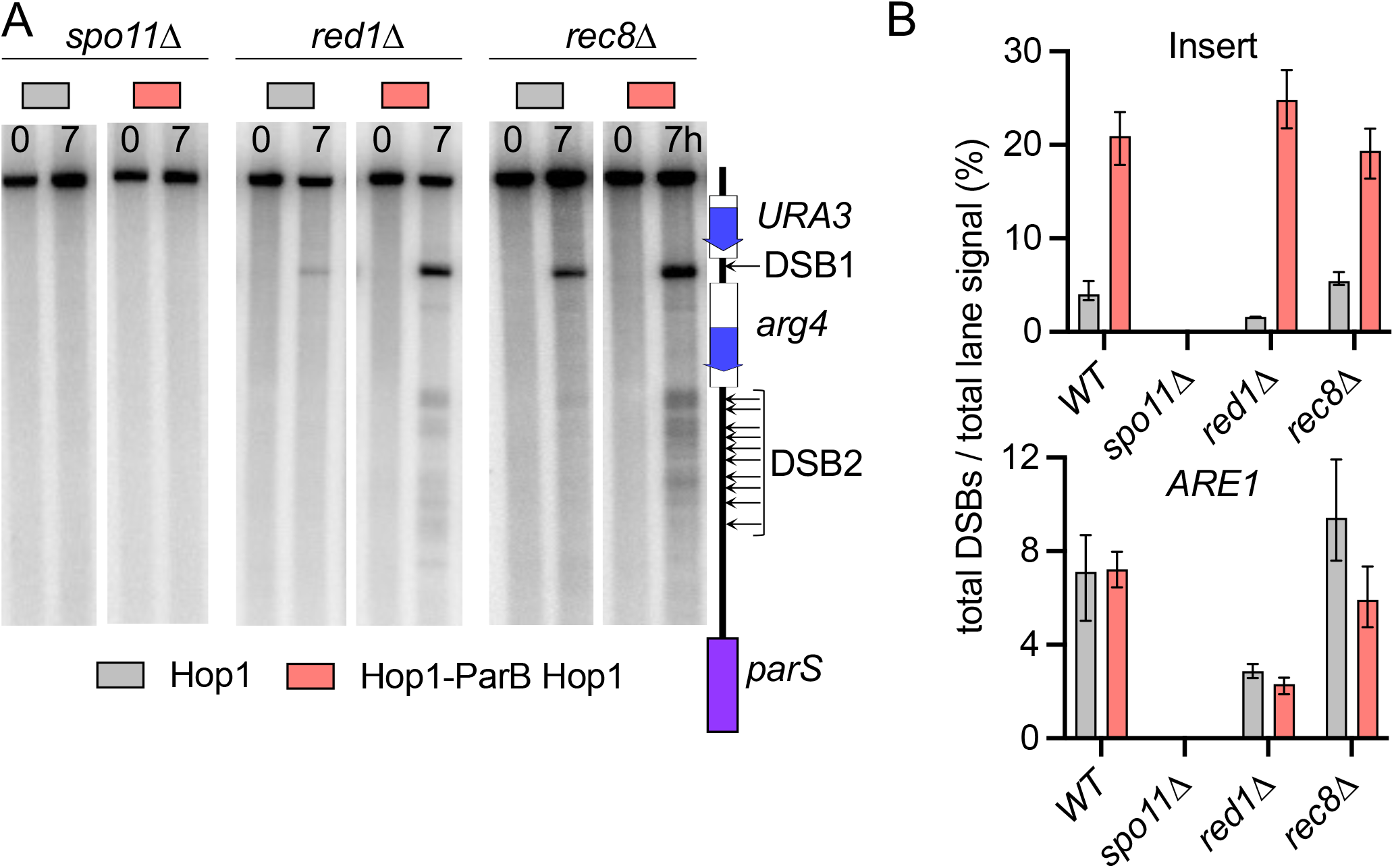
Hop1-stimulated DSBs are Spo11 dependent but Red1- and Rec8-independent. A. Southern blot with DNA from a *sae2Δ* strain with inserts at *URA3* in *spo11Δ, red1Δ* or *rec8Δ* strains homozygous either for *HOP1* (grey) or *HOP1-parB HOP1* (salmon). DNA was digested with *Sbf*I and probed with *parS* sequences. B. Top—DSBs in the *parS* insert at *URA3*, measured at 7h after induction of meiosis. Bottom—DSBs in the same strains at the *ARE1* control locus. See also File S1 sheet 5.

Previous studies have show that *rec8Δ* strains display rearranged patterns of Red1, Hop1 and Spo11-complex components, with a tendency towards reducing occupancy at hotspots in the centers of large chromosomes while preserving occupancy, albeit at much-reduced levels, on short chromosomes and at certain loci on other chromosomes (Kugou *et al*. 2009; Panizza *et al*. 2011; Sun *et al*. 2015; Heldrich *et al*. 2022). Consistent with previous data showing that occupancy by these proteins is not substantially altered in *rec8Δ* mutants at *URA3* and *ARE1* (Kugou *et al*. 2009; Panizza *et al*. 2011), DSBs at the *parS* insert locus and at the *ARE1* control locus were similar in *REC8* and in *rec8Δ* strains, regardless of the presence or absence of Hop1-ParB (Figures 4A and 4B; File S1 sheet 5). Thus, unlike many DSB hotspots in the centers of long chromosomes, the hotspot created by recruitment of Hop1-ParB to the insert at *URA3* is not affected by loss of Rec8-cohesin.

### The ParB-parS system specifically enriches Hop1 at the target locus

To confirm that the ParB/*parS*-dependent increase in meiotic recombination was associated with recruitment of Hop1, we used calibrated ChIP-seq to map Hop1 occupancy genome wide, using a spike-in sample from meiotic cells of *Saccharomyces mikatae. S. mikatae* is substantially diverged from *S. cerevisiae* (24% nucleotide divergence genome wide), but *S*.*mikatae* Hop1 shows 86.5% amino acid identity with *S. cerevisiae* Hop1, and cross-reacts with the antiserum against *S. cerevisiae* Hop1 used here for ChIP ((Kellis *et al*. 2003; Dujon 2006; Liti *et al*. 2013; Lam and Keeney 2015); data not shown). Strains expressing either both Hop1 and Hop1-ParB (*HOP1-ParB HOP1/HOP1*) or Hop1 alone displayed similar Hop1 occupancy profiles genome-wide. However, Hop1 occupancy in a ∼50 kb region surrounding the *parS* insert at *URA3* was much greater than the Hop1 signal in the rest of the genome (Figures 5A and 5C). Quantitative interpretation of this pattern is complicated by the fact that strains expressing both Hop1-ParB and Hop1 have three modes of Hop1 chromosome binding: direct binding of the ParB domain in Hop1-ParB to chromosomal DNA; indirect binding of Hop1 through its interactions with itself and with Red1 and cohesin (Smith and Roeder 1997; Klein *et al*. 1999; Blat *et al*. 2002; Riedel *et al*. 2006; Panizza *et al*. 2011; Sun *et al*. 2015; West *et al*. 2018; West *et al*. 2019); and possible direct binding of Hop1 to DNA (Kironmai *et al*. 1998; Kshirsagar *et al*. 2017; Heldrich *et al*. 2020; Heldrich *et al*. 2022). Hop1 bound in these three modes is likely to be crosslinked to DNA with different efficiencies. Therefore, while the increased Hop1 ChIP signal in the vicinity of *parS* almost certainly indicates that more Hop1 is bound in this region, the quantitative extent of that increase remains to be determined.

**Figure 5:**
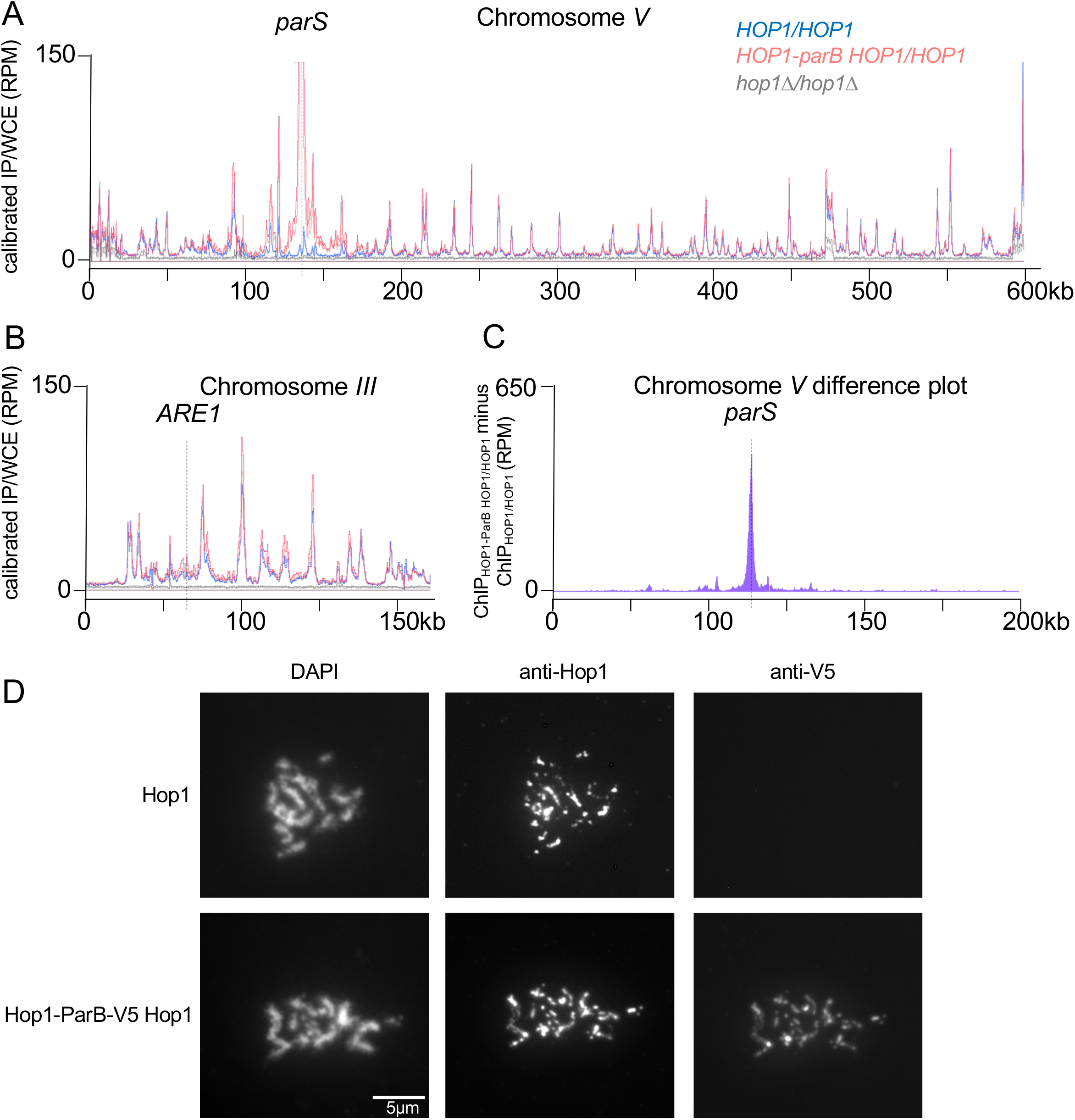
Hop1 localization and enrichment at *parS*. A. Hop1 occupancy (immunoprecipitate/whole cell extract) on chromosome *V*, determined by calibrated ChIP-seq (see Materials and Methods) using samples taken at 4h after induction of meiosis. Strains contained the *URA3-tel-arg4-parB* insert at *URA3* and the indicated *HOP1* genotype. Dark and light lines indicate replicates. Dotted vertical line—*parS* insert locus. The peak at *parS* in *HOP1-parB HOP1/HOP1* strains is truncated; peak values reached ∼700 RPM. B. Hop1 occupancy around the *ARE1* control locus (chr. *III*). Dotted vertical line—*ARE1* DSB site. All other details as in A. C. Difference plot for 200 kb around *parS*, calculated by subtracting the calibrated ChIP/WCE for *HOP1/HOP1*(mean of both replicates) from that for *HOP1-parB HOP1*/*HOP1* (mean of both replicates). D. Chromosome spreads from meiotic cells (4 and 5 h) from wild-type and from cells expressing Hop1-ParB (*HOP1-parB-V5 HOP1/HOP1*), probed with the indicated antiserum. In strains expressing Hop1-parB-V5, Hop1-ParB (anti-V5) shows the same distribution as total Hop1. Scale bar =5µm

We also compared the distribution of Hop1 and Hop1-ParB on chromosome spreads of cells at the pachytene stage, using anti-Hop1 to detect both proteins and anti-V5 to specifically detect the Hop1-ParB fusion protein (Figure 5C). Cells expressing Hop1 alone displayed a pattern of lines and punctate foci, as has been previously reported (Smith and Roeder 1997). Cells expressing both Hop1 and Hop1-ParB displayed a similar pattern, and similar staining patterns were obtained with anti-Hop1 (detecting Hop1 and Hop1-ParB) and anti-V5 (detecting Hop1-ParB only). Thus, Hop1-ParB appears to localize across the genome.

### Hop1-ParB provides partial Hop1 function

The experiments described above used strains with a wild-type copy of the *HOP1* gene in addition to the *HOP1-ParB* fusion (see Materials and Methods). To determine whether Hop1-ParB was fully functional when expressed on its own, we examined meiotic spore viability, recombination, and DSB formation in strains containing a *parS* insert at *URA3* where the only source of Hop1 was a Hop1-ParB fusion (Figure 6). In strains where only Hop1-ParB was expressed (*HOP1-PARB/HOP1-ParB*), spore viability was reduced ∼2-fold (Figure 6A; File S1 sheet 7), recombination between *arg4* heteroalleles inserted at *URA3* was reduced ∼3-fold (Figure 6B; File S1 sheet 4), and DSBs at the *parS* insert locus and at the *ARE1* control locus were reduced ∼3 and ∼6-fold, respectively (Figures 6C and 6D; File S1 sheet 5), relative to strains expressing both Hop1 and Hop1-ParB (*HOP1-ParB HOP1/HOP1-ParB HOP1*). These defects were at least partially suppressed by the addition of a single copy of untagged *HOP1* (*HOP1-ParB/HOP1*), while full suppression of the DSB defect required two wild-type copies of Hop1. Thus, while the Hop1-ParB fusion construct produces a protein that can recruit Hop1 protein to the region surrounding *parS*, it is unable to provide full Hop1 function.

**Figure 6.**
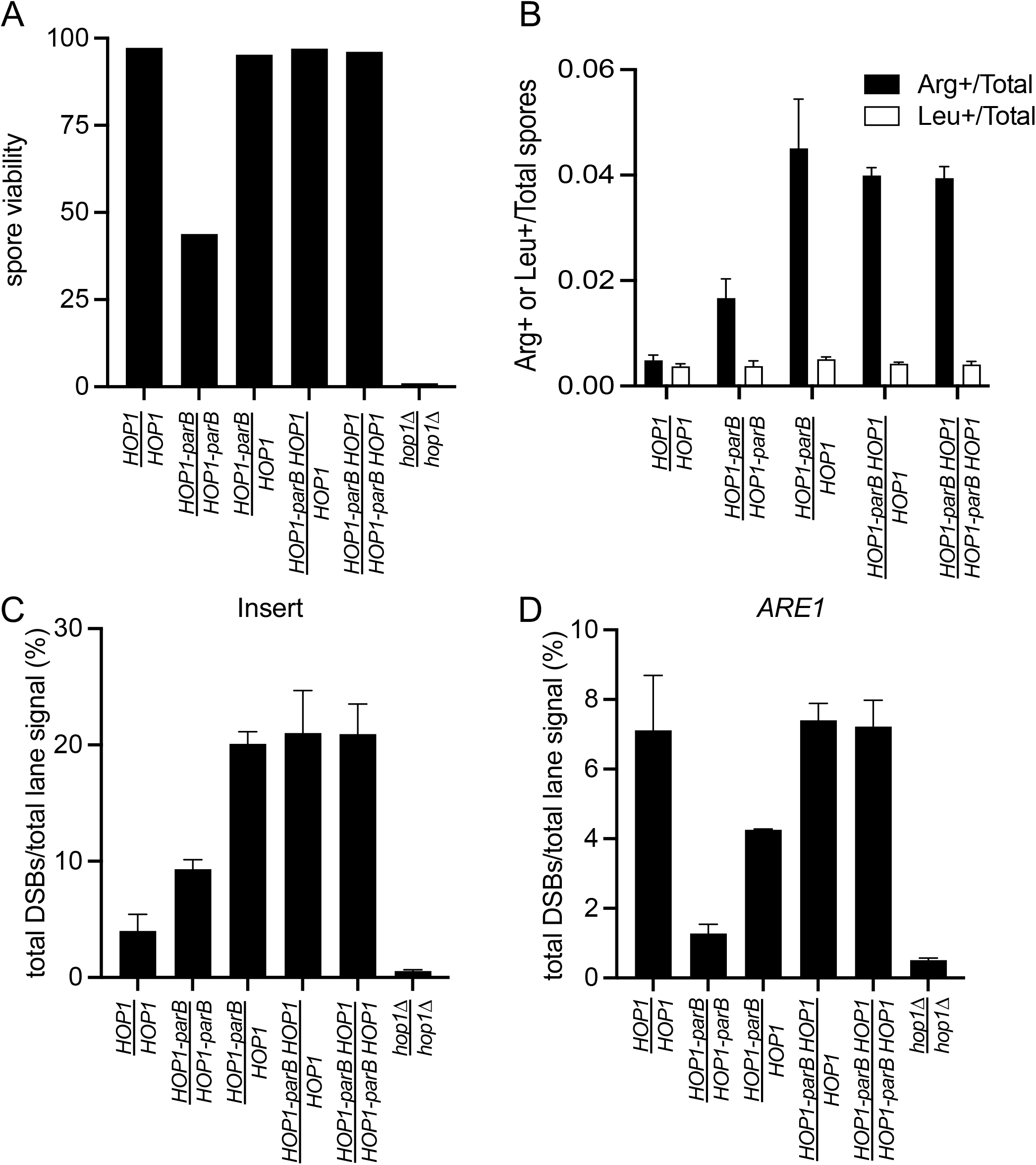
Hop1-ParB has partial function. A. Spore viability in dissected tetrads. B. Frequencies of Arg+ (black) and Leu2+ (white) recombinants in random spores from diploids with the *URA3-arg4-parS* insert at *URA3*. C, D. Frequencies of DSBs (DSB1 + DSB2, see Figure 3) at the *URA3-arg4-parS* insert at *URA3* and at the *ARE1* control locus, in *sae2Δ* strains. See also File S1 sheets 4, 5, and 7.

### Non-canonical DSB repair in the presence of Hop1-ParB and in inserts at URA3

The Hop1-ParB fusion protein also conferred an apparent defect in meiotic DSB repair. Cells expressing Hop1-ParB showed a 45-50 min delay in the disappearance of DSBs at both the *parS* insert and at *ARE1* (Figures 7A,C,D; File S1 sheet 8). This delay was accompanied by a delay in meiotic divisions that increased with *HOP1-parB* gene dosage (Figure 7B; File S1 sheet 9), consistent with the presence of unrepaired DSBs activating the meiotic checkpoint (Lydall *et al*. 1996; Grushcow *et al*. 1999; Thompson and Stahl 1999; Roeder and Bailis 2000; Shimada *et al*. 2002).

**Figure 7.**
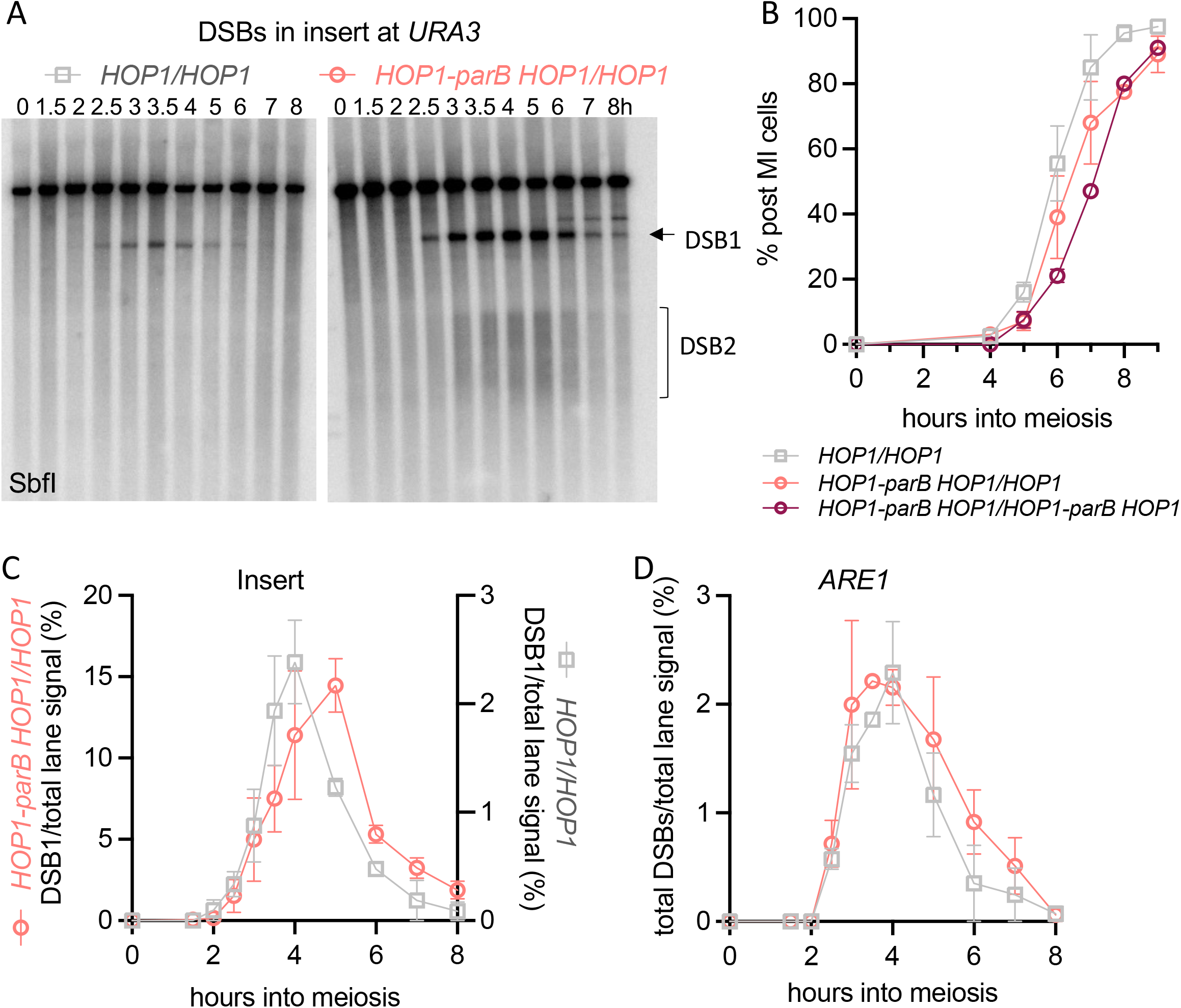
Delayed DSB repair and meiotic progression in presence of Hop1-ParB. A. Southern blots of meiotic DNA from *SAE2* cells with the *URA3-arg4-parS* insert at *URA3*, expressing either Hop1 or both Hop1-ParB and Hop1 digested with *Sbf1* and probed with *parS* sequences. B. Meiotic progression, expressed as cells completing meiosis I (with either 2 or 4 nuclei). C. Quantification of total DSBs from the experiment in panel A and others. Note that data from strains expressing Hop1 (grey) are plotted with the right-hand Y axis, and from strains with *HOP1-ParB HOP1/HOP1* (salmon) on the left-hand Y axis. D. DSBs at the *ARE1* control locus from the same experiments. See also File S1 sheets 8, 9.

In addition, Hop1-ParB-stimulated crossovers in inserts at *URA3* do not appear to use the recombination pathway that is dominant at other DSB hotspots. Previous studies indicate that most meiotic crossovers form via a pathway that involves the ZMM protein complex and the meiosis-specific MutLγ resolvase (Mlh1-Mlh3-Exo1), and a minor fraction are formed by structure-selective nucleases (SSNs; Mus81-Mms4, Yen1, Slx1-Slx4) (Schwacha and Kleckner 1994; Wang *et al*. 1999; Khazanehdari and Borts 2000; Kirkpatrick *et al*. 2000; Tsubouchi and Ogawa 2000; Allers and Lichten 2001a; Allers and Lichten 2001b; De los Santos *et al*. 2003; Hoffmann *et al*. 2003; Argueso *et al*. 2004; Bishop and Zickler 2004; Borner *et al*. 2004; Hollingsworth and Brill 2004; Jessop *et al*. 2006; Lynn *et al*. 2007; Jessop and Lichten 2008; Nishant *et al*. 2008; Zakharyevich *et al*. 2010; Comeron *et al*. 2012; De Muyt *et al*. 2012; Wang *et al*. 2012; Yang *et al*. 2012; Zakharyevich *et al*. 2012; Agostinho *et al*. 2013; Oke *et al*. 2014; Al-Sweel *et al*. 2017; De Muyt *et al*. 2018; Pyatnitskaya *et al*. 2019). However, genetic crossing-over in a ∼25 kb interval containing the *URA3*-*arg4*-pBR322-*parS* was reduced only modestly in *mlh3Δ* strains, both in strains where Hop1-ParB was expressed and where Hop1-parB was absent (Figure 8; File S1 sheet 10).

**Figure 8.**
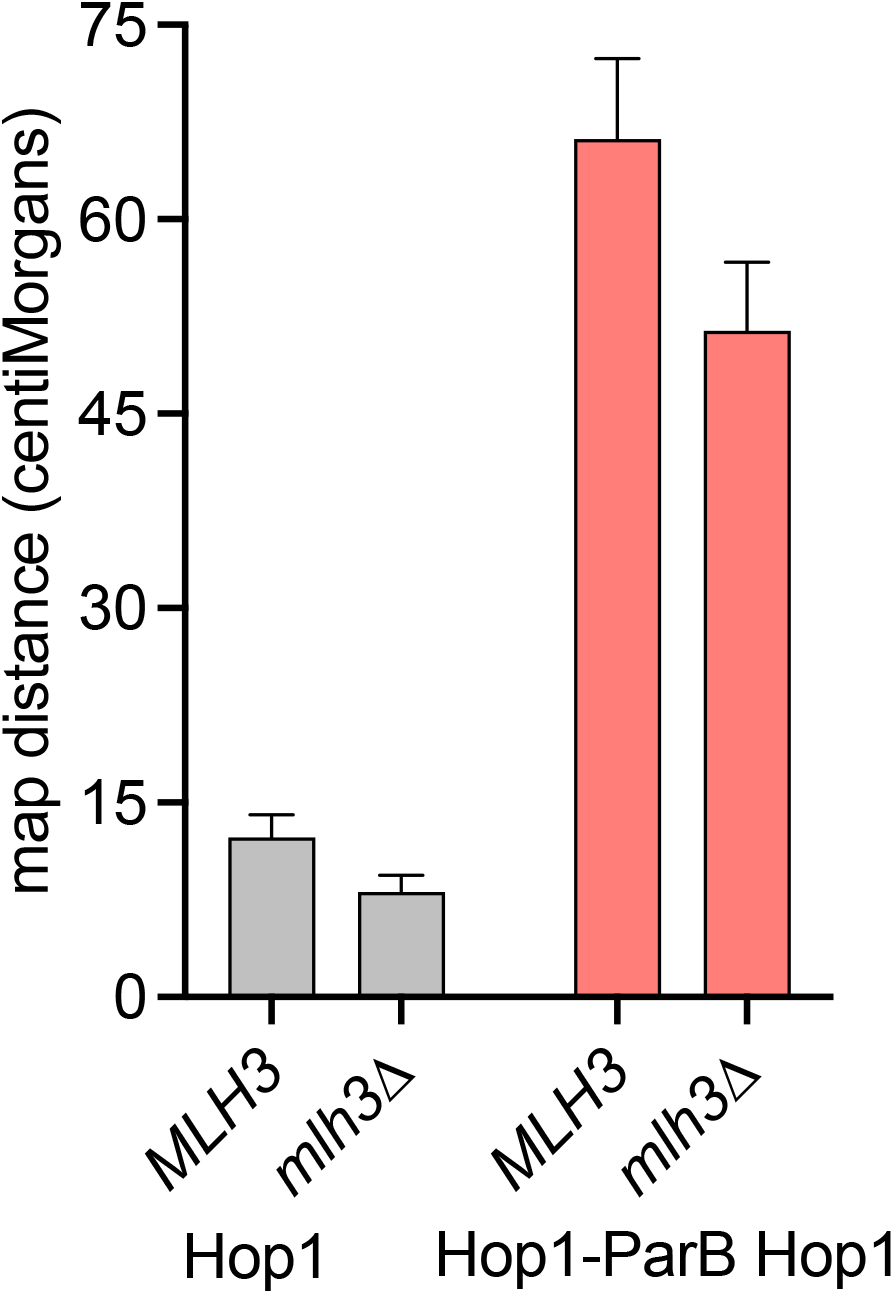
Most crossovers at *URA3* are MutLγ -independent. Map distances, calculated from marker segregation in tetrads, between *natMX* and *hygMX* inserts flanking a *URA3-arg4-parS* insert at the *URA3* locus (see Materials and Methods). Expression of Hop1-ParB results in a marked increase in map distances. Map distances are only modestly decreased in *mlh3Δ* strains, in either *HOP1/HOP1* strains (grey) or *HOP1-parB HOP1/HOP1-parB HOP1* strains (salmon). See also File S1 sheet 10.

We also measured crossover (CO) and noncrossover (NCO) recombination at the molecular level, using a second *parS-*containing insert at *URA3* (*URA3-tel-ARG4-parB*; Figure 9A) that contains a single DSB site (Jessop *et al*. 2005; Ahuja *et al*. 2021). Consistent with experiments described above that used *URA3-arg4-*pBR322-*parS* inserts, addition of a single copy of Hop1-ParB (*HOP1-parB HOP1/HOP1*) resulted in a marked increase in DSBs (∼7-fold; Figures 9A and 9B; File S1 sheet 12), in COs (∼5-fold; Figure 9A and 9C; File S1 sheet 11), and in NCOs (∼4.5-fold; Figure 9A and 9D; File S1 sheet 11) within the insert at *URA3*.

**Figure 9.**
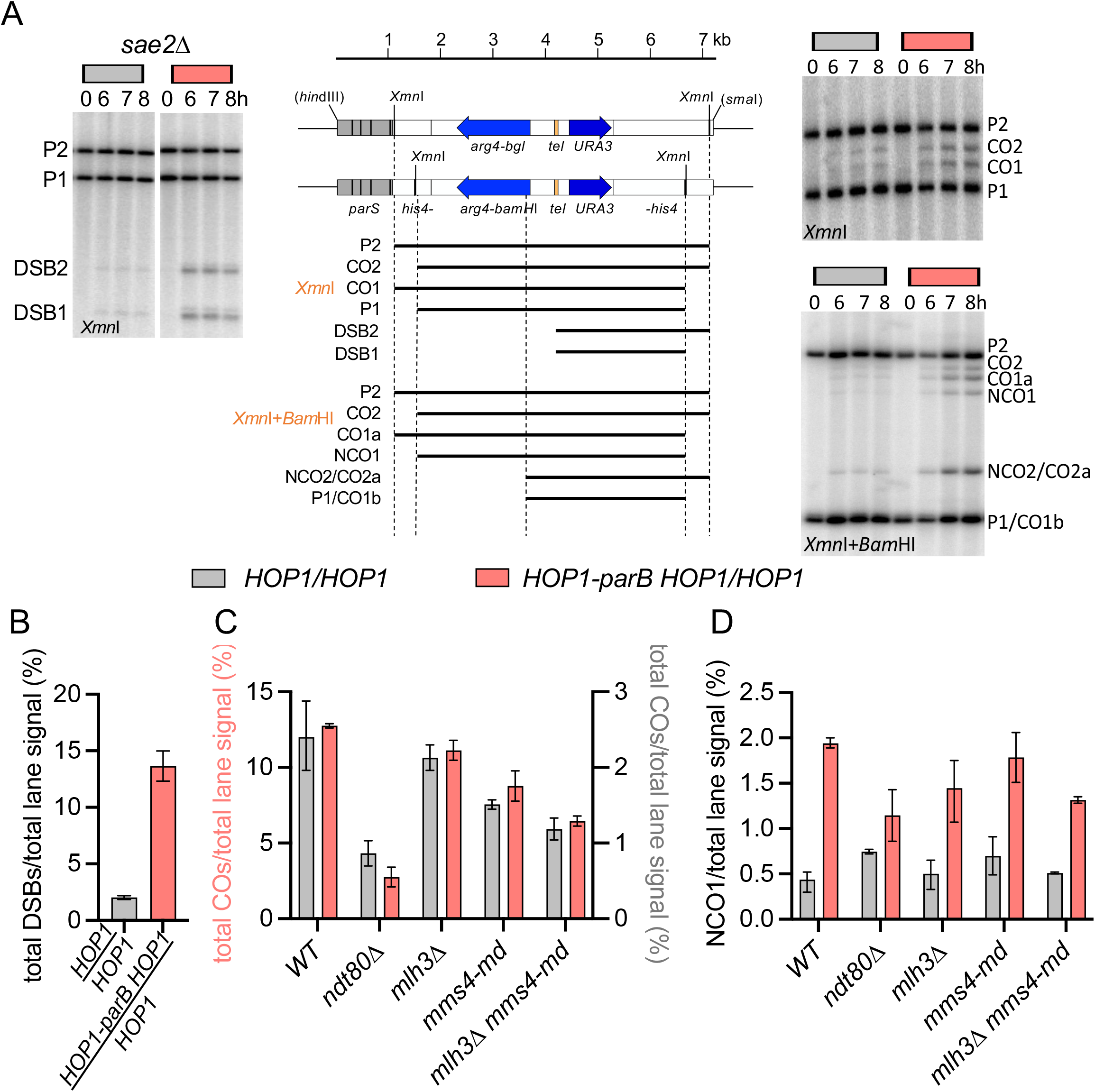
Noncanonical crossover pathway usage at *URA3*. A. Schematic for the *URA3-tel-arg4-parS* reporter insert at *URA3*, showing product length in *Xmn*I or *XmnI+Bam*HI digests. Left—Southern blots showing DSBs in *sae2Δ* strains; right—recombination products in *SAE2* strains. Top—crossovers (CO1 and CO2) in *Xmn*I; bottom—noncrossovers (NCO1) in *Xmn*I*+Bam*HI digests. Both blots were probed with *URA3* sequence. B. Quantification of insert DSBs in *sae2Δ HOP1/HOP1*(grey) or *sae2Δ HOP1-parB HOP1/HOP1* (salmon) strains in samples taken 7h after meiotic induction. C. Quantification of COs (CO1 + CO2) in *HOP1/HOP1* (grey, right Y axis) or *HOP1-ParB HOP1/HOP1* (salmon, left Y axis) in samples taken 8h after meiotic induction in the indicated mutants. D. Quantification of NCOs (NCO1). Details as in C. See also File S1 sheets 11, 12.

Previous studies have shown that *ndt80Δ* mutants arrest at the pachytene stage of meiosis with markedly reduced CO levels, regardless of whether MutLγ or SSNs are the primary resolvase (Xu *et al*. 1995; Chu and Herskowitz 1998; Allers and Lichten 2001a; Jessop and Lichten 2008; Sourirajan and Lichten 2008; De Muyt *et al*. 2012). Consistent with this, COs in the insert at *URA3* were reduced 3 to 4-fold in *ndt80Δ* mutants, regardless of whether Hop1-ParB was present or absent (Figure 9C; File S1 sheet 11). However, unlike at other hotspots, where *mlh3Δ* causes a ∼2-fold reduction in COs (Hunter and Borts 1997; Wang *et al*. 1999; Argueso *et al*. 2004; Nishant *et al*. 2008; Al-Sweel *et al*. 2017), *mlh3Δ* caused a much more modest 10-15% decrease in COs (Figures 9C; File S1 sheet 11), as was seen in genetic crosses (Figure 8).

Inactivation of the major mitotic resolvase Mus81-Mms4, in *mms4-md* mutants, reduced COs in inserts at *URA3* by 30-40% (Figure 9C; File S1 sheet 11), as compared to reductions of 10-20% (*mus81* or *mms4*) seen at recombination hotspots (Borner *et al*. 2004; Jessop *et al*. 2006; Oh *et al*. 2007; De Muyt *et al*. 2012; Zakharyevich *et al*. 2012; He *et al*. 2020). Furthermore, COs were reduced about 2-fold in *mlh3 mms4-md* double mutants, as compared to >6-fold reductions reported in other genetic studies (Argueso *et al*. 2004; Nishant *et al*. 2008; Brown *et al*. 2013). Notably, COs were similarly affected in mutant strains whether Hop1-ParB was present or absent, indicating that the non-canonical CO pathway utilization seen in inserts at *URA3* is independent of DSB levels.

## DISCUSSION

The meiotic chromosome axis lies at the center of meiotic chromosome transactions, including the initiation of recombination by double strand break formation, recombination partner choice and homolog pairing, CO designation and pathway choice, and CO assurance and spacing control (Hollingsworth and Ponte 1997; Zickler and Kleckner 1999; Blat *et al*. 2002; Glynn *et al*. 2004; Kleckner 2006; Tsubouchi and Roeder 2006; Niu *et al*. 2007; Yang *et al*. 2008; Joshi *et al*. 2009; Kugou *et al*. 2009; Niu *et al*. 2009; Callender and Hollingsworth 2010; Kim *et al*. 2010; Panizza *et al*. 2011; Youds and Boulton 2011; Chuang *et al*. 2012; De Muyt *et al*. 2018; Pyatnitskaya *et al*. 2019; Ur and Corbett 2021). While axis proteins’ roles in these processes have been extensively studied, the co-dependent localization of axis proteins has presented a challenge to the identification of their individual roles in meiotic recombination. In this paper, we used the bacterial ParB-*parS* system to independently enrich the axis protein Hop1 at target loci, and to identify a unique role for Hop1 in DSB formation.

### Hop1 is efficiently recruited by the ParB-parS system

While the ParB-*parS* system has previously been used as an alternative to operator-repressor arrays to visually label specific loci (Saad *et al*. 2014; Germier *et al*. 2018), the use of this system to recruit meiotic axis proteins is, to our knowledge, the first time that it has been used to localize chromosomal proteins with the goal of understanding their function. Hop1-ParB expression caused a markedly greater increase in Hop1 ChIP signal at the *parS* site and for about 25 kb to either side of *parS*, consistent with the spread of ParB from *parS* observed in bacteria, which is facilitated by its ability to dimerize and form a clamp that slides along DNA (Walter *et al*. 2020) and by its ability to bridge DNA (Breier and Grossman 2007; Graham *et al*. 2014; Antar *et al*. 2021). Cytological analysis showed that Hop1-ParB and the wild-type Hop1 protein show similar nucleus-wide localization patterns (Figure 5D), suggesting that the C-terminaltag does not prevent Hop1-ParB loading via Hop1-Red1 or Hop1-Hop1 interactions (West *et al*. 2018; West *et al*. 2019).

### Hop1 determines local DSB levels

Previous studies have reported a direct correlation between levels of Hop1 (and Red1) enrichment and levels of Spo11 DSBs in different regions of the genome (Pan *et al*. 2011; Panizza *et al*. 2011; Subramanian *et al*. 2019). Here, we have shown that ParB-*parS-*mediated recruitment of Hop1 to a locus causes a dramatic increase in Spo11-dependent DSBs at that locus. This increase in DSBs is independent of the other meiosis-specific axis proteins, Red1 and Rec8. Thus, while Red1 and Rec8 might be required for Hop1 loading under normal circumstances, it is the level of Hop1 enrichment that ultimately determines the local DSB levels. This suggests that Hop1 alone is sufficient to recruit the DSB-forming Spo11 complex consistent with recent biochemical studies showing that the Spo11 complex protein Mer2 interacts directly with Hop1 and not with Red1 (Rousova *et al*. 2021).

We found that, while Hop1-ParB can stimulate DSB formation in the vicinity of *parS*, Hop1-ParB alone was insufficient for optimal DSB formation, recombination, and spore viability, and that full function required addition of 1 to 2 copies of wild-type *HOP1*, depending upon the assay (Figure 6). Since Hop1-ParB is produced at about 80-90% of the levels of wild-type Hop1 protein (Figure 1B), it is unlikely that these results can be explained by reduced levels of Hop1 protein alone, although it is possible that over-enrichment of Hop1 at *parS* reduces Hop1 levels elsewhere in the genome. It also is possible that the presence of the C-terminal ParB tag creates a partially functional Hop1 protein. For example, recent *in vitro* studies have shown Mer2 preferentially binds to Hop1 with an unlocked closure motif (Rousova *et al*. 2021). Chromosome-bound Hop1 normally is in this unlocked configuration, due to closure motif-HORMA domain interactions that recruit it to the axis(West *et al*. 2018; West *et al*. 2019). However, Hop1 recruited to chromosomes by a ParB tag might frequently exist in the locked confirmation, and thus might recruit the Spo11 complex less efficiently. In addition, the ParB tag might interfere with interactions necessary for Hop1 post-translational modification, and/or Hop1 loading/unloading (Carballo *et al*. 2008; Wojtasz *et al*. 2009; Thacker *et al*. 2014; Herruzo *et al*. 2016; Herruzo *et al*. 2021; Li and Shinohara 2021). For example, Hop1 is normally removed from the axis after homolog synapsis (Borner *et al*. 2008; Joshi *et al*. 2009; Wojtasz *et al*. 2009; Zanders and Alani 2009; Daniel *et al*. 2011; Kauppi *et al*. 2013; Thacker *et al*. 2014; Lambing *et al*. 2015; Subramanian *et al*. 2016; Subramanian *et al*. 2019); a failure to remove Hop1-ParB bound via the ParB tag might result in the persistent DSBs and delayed progression that we observed when Hop1-ParB is present (Figure 7).

### Noncanonical recombination pathway usage in inserts at the URA3 locus

Previous studies of crossover formation have concluded that most meiotic COs are formed by MutLγ -dependent double Holliday junction resolution, a minor fraction are formed by mitotic resolvases (Mus81-Mms4/Eme1 and Yen1/Gen1) and that both modes of resolution are *CDC5*- and *NDT80*-dependent (Xu *et al*. 1995; Chu and Herskowitz 1998; Allers and Lichten 2001a; Clyne *et al*. 2003; Jessop and Lichten 2008; Sourirajan and Lichten 2008; De Muyt *et al*. 2012; Schwartz *et al*. 2012; Zakharyevich *et al*. 2012; Blanco and Matos 2015; Yoon *et al*. 2016). These studies either examined events at recombination hotspots or examined crossing-over in large genetic intervals, in which presumably most recombination is hotspot-driven. We find that recombination in inserts at *URA3* does not conform to these conclusions. While *ndt80Δ* substantially reduced COs (Figure 9C), consistent with crossing over in inserts at *URA3* being resolvase-driven, specific resolvase-dependence of COs was substantially altered. Unlike in previous studies, where loss of MutLγ results in a ∼2-fold CO reduction (Hunter and Borts 1997; Wang *et al*. 1999; Argueso *et al*. 2004; Nishant *et al*. 2008; Al-Sweel *et al*. 2017), *mlh3Δ* mutants showed a substantially lower CO reduction (∼ 20-25% when measured genetically, Figure 8; ∼10 to 15% at the molecular level, Figure 9C). In addition, *mms4-md* mutants, which cause a meiosis-specific loss of Mus81-Mms4 activity, showed a substantial (30-40%) CO reduction in inserts at *URA3*, which is greater than the minor CO reductions seen in the absence of Mus81-Mms4 in other studies (Argueso *et al*. 2004; Jessop and Lichten 2008; De Muyt *et al*. 2012; Zakharyevich *et al*. 2012). Taken together, these data indicate a shift away from resolution by MutLγ, and towards resolution by mitotic resolvases during Spo11-induced recombination at *URA3*, as was seen for recombination during meiosis that is initiated by the VDE site-specific endonuclease (Medhi *et al*. 2016; Shodhan *et al*. 2019). Of particular importance, similar resolvase-usage patterns are seen in *HOP1/HOP1* and *HOP1-parB HOP1/HOP1* strains, even though DSB levels and CO levels differ more than 5-fold between these strains (Figures 9B and 9C). Thus, non-physiological recruitment of Hop1 appears to have created a DSB hotspot that still behaves like a coldspot during the post-DSB steps of recombination.

One possible explanation for this is that Hop1-independent features of chromosome structure determine CO pathway choice. One such feature could be the axis proteins themselves. Red1 interacts with Zip4, a part of the ZZS complex and the larger ZMM protein complex that is important for the MutLγ -dependent CO pathway, and this interaction is conserved in other organisms (Yang *et al*. 2008; De Muyt *et al*. 2018; Pyatnitskaya *et al*. 2019). The meiotic cohesin component Rec8 has also been identified as to playing a role in homolog bias and CO formation (Yoon *et al*. 2016; Hong *et al*. 2019), although this may simply reflect its role in recruiting Red1. If recruiting Hop1 to “cold” regions increases DSB formation without increasing meiotic cohesin and/or Red1 levels, it is possible that insufficient ZMM proteins are recruited to promote MutLγ -dependent intermediate resolution, leading to an increased use of mitotic resolvases during CO formation.

In summary, we report here the novel use of the ParB-ParS bacterial partition system, to study the role of chromosome structural proteins in meiotic recombination at a specific locus without substantially altering recombination elsewhere in the genome.

The artificial recruitment of Hop1 to regions where meiotic axis proteins are normally low enabled the conversion of DSB/recombination coldspots into recombination hotspots by specifically increasing DSB formation independent of other axis proteins. Our data suggest an independent role for Hop1 in DSB formation, but also a need for the other axis proteins or other factors in normal repair pathway choice. It will be of interest to determine if recruiting both Red1 and Hop1 to these coldspots-turned-hotspots can restore a more wild-type pattern of resolvase usage during meiotic CO formation. We also anticipate that artificial Hop1 recruitment could facilitate analysis of the interactions between Hop1 and Spo11-complex proteins that promote DSB formation. In addition, artificial recruitment of Hop1 homologs in other organism may provide a targeted way to increase meiotic recombination in regions where recombination is normally low, both for mechanistic studies and for breeding purposes.

## Supporting information

File S1-underlying data

File S2-ChIP calibration script

File S3-pMJ1088 sequence (genbank)

## DATA AVAILABILITY

All strains are available upon request. The authors affirm that all data necessary to confirm the conclusions of this article are represented within the article, tables, figures, supplementary materials, and publicly available database depositions. Numerical values underlying graphs in all Figures are provided in File S1; sequence reads are available at GEO (https://www.ncbi.nlm.nih.gov/geo/, accession number GSE201240).

## ACKNOWLEDGEMENTS

We thank Darpan Medhi for suggesting the use of ParB-*parS* to recruit axis protein, Jasvinder Ahuja, Matan Cohen, Julie Cooper, Kevin Corbett, Yolanda L. Jones, and Alex Kelly for discussions and comments on the manuscript, Kevin Corbett for the gift of Hop1 protein used for antisera, and the Center for Cancer Research Genomics Core for high-throughput sequencing. This work used resources at the NIH HPC Biowulf cluster (https://hpc.nih.gov) and is supported by the Intramural Research Program of the NIH through the Center for Cancer Research at the National Cancer Institute. The funders had no role in research design, execution, analysis, interpretation or reporting. The authors have no conflicting interests to report.

## FIGURE LEGENDS

**Figure S1.**
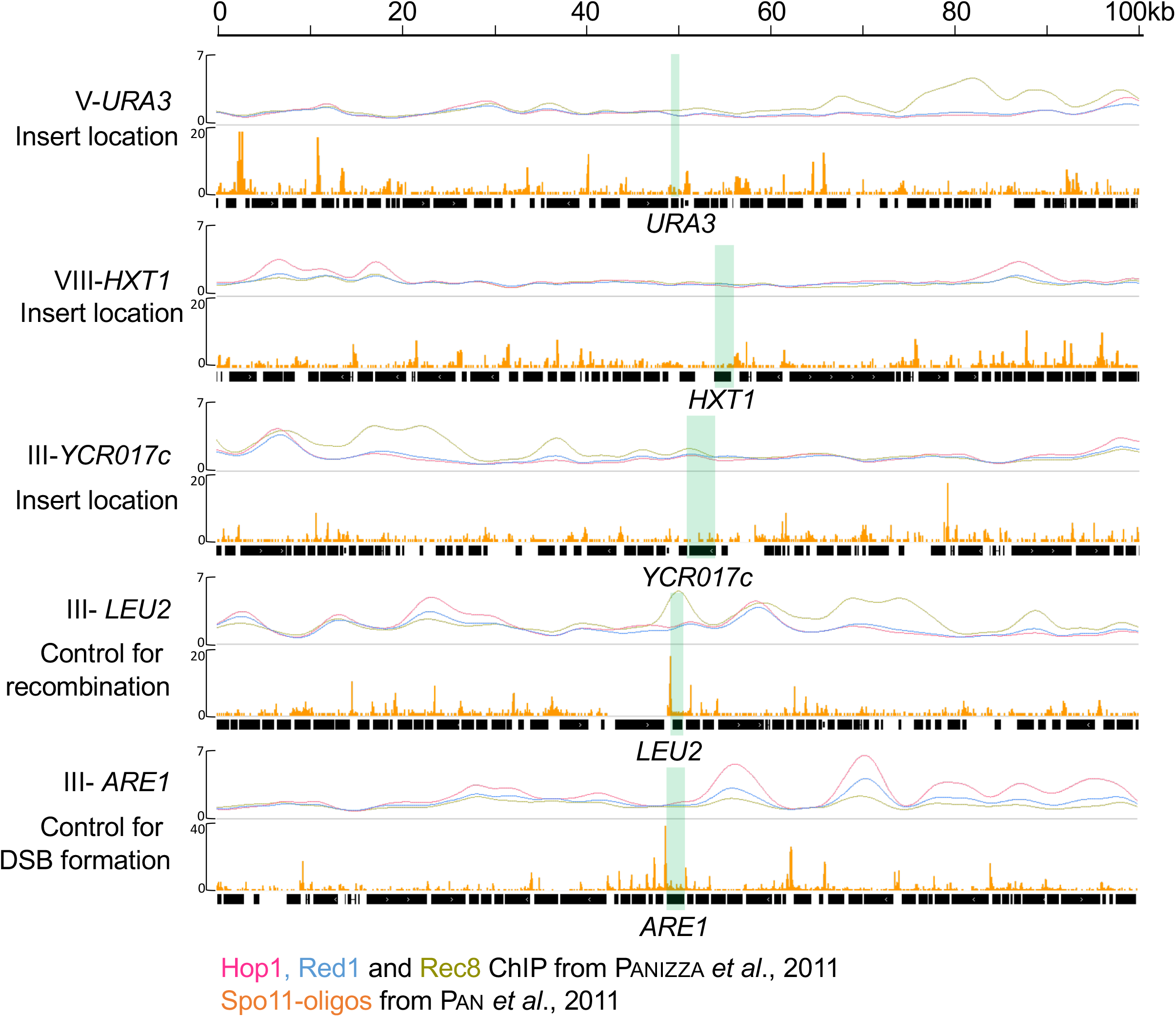
Genome-wide axis protein localization. Line plots—levels of Hop1 (pink), Red1 (blue) and Rec8 (olive green), at the locations of the three inserts and at the two control loci, expressed as decile-normalized ChIP/WCE, data from Panizza *et al*. (2011). Orange bars-–levels of Spo11 DSBs, counts of Spo11-linked oligonucleotides (hPM/bp), data from Pan *et al*. (2011). Green boxes—gene positions.

**Figure S2.**
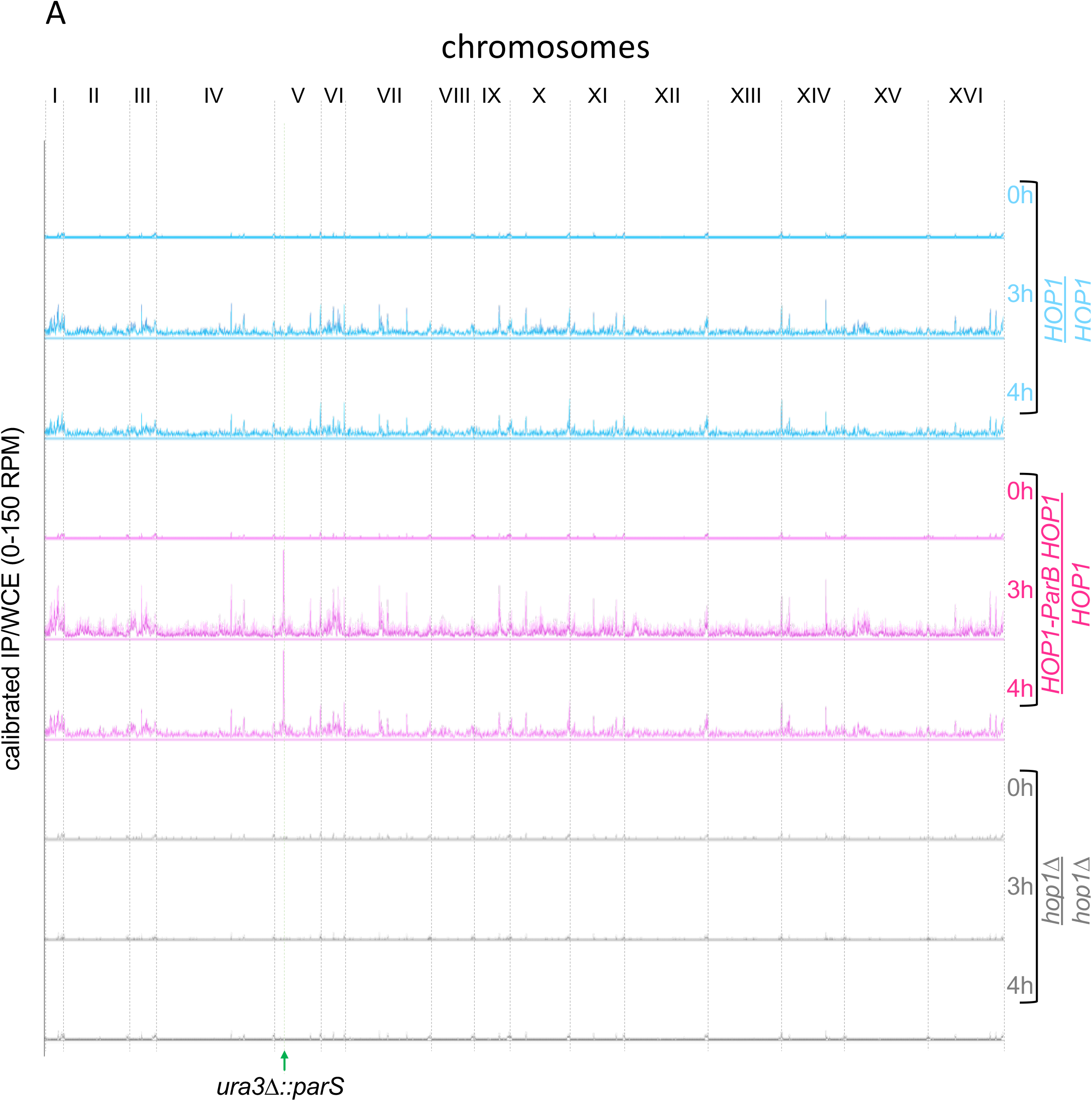
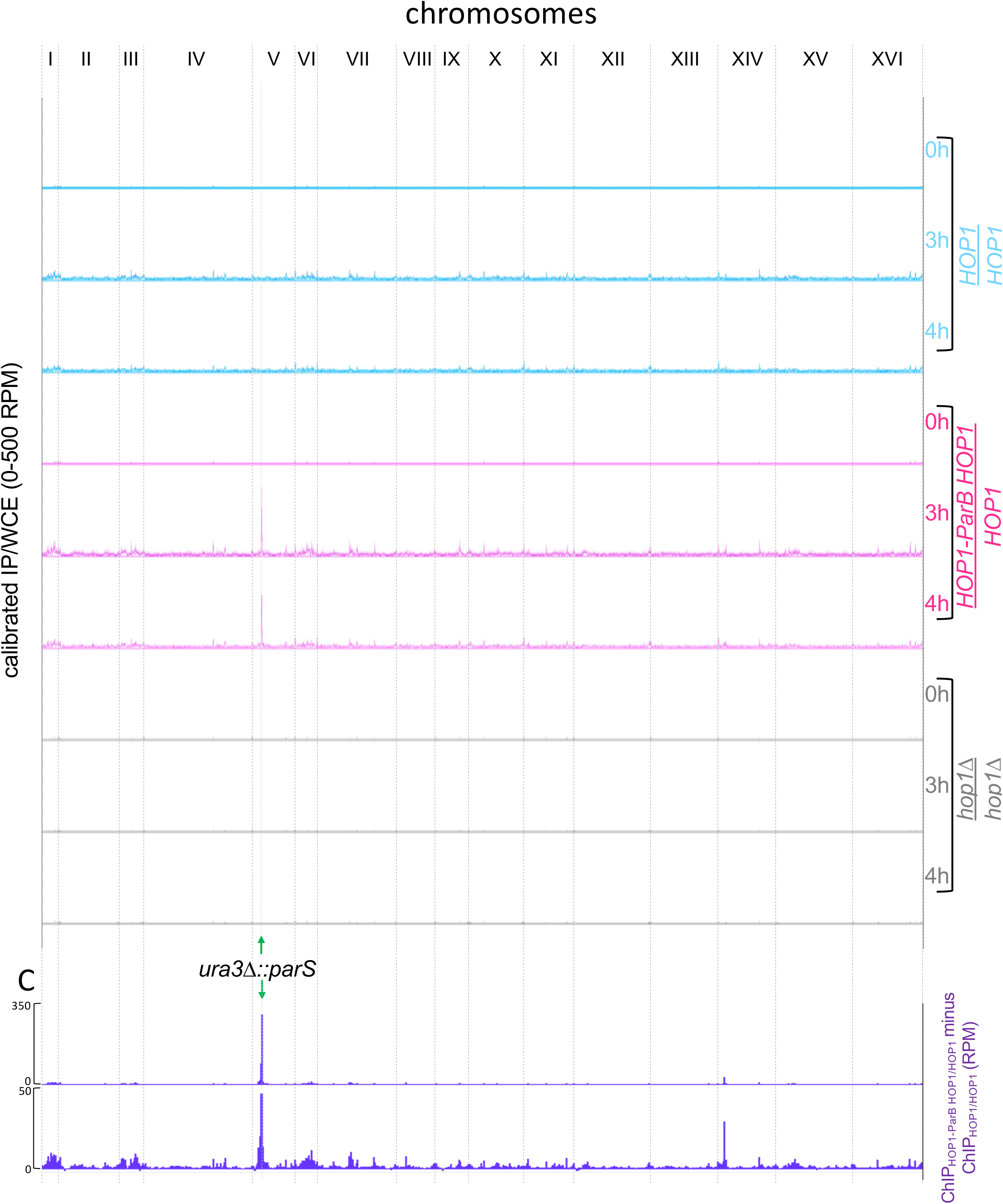
Genome-wide Hop1 localization. Whole-genome Hop1 occupancy (immunoprecipitate/whole cell extract) determined by calibrated ChIP-seq of samples taken at 0h, 3h and 4h in meiosis from strains with a *URA3-tel-arg4-parB* insert at *URA3* and the indicated *HOP1* genotype, plotted using a bin size of 7kb. Vertical grey lines—chromosome boundaries. Vertical green line—*parS* insert location (chr. *V*). The two replicates are indicated by dark and light lines. A. Y axis scale of 0-150 RPM to illustrate Hop1 occupancy across the whole genome; the signal peak at the *parS* insert location. B. As in panel A, but with Y axis scale of 0-500 RPM to illustrate the full range of Hop1 occupancy. C. Difference plot calculated by subtracting the calibrated ChIP/WCE for Hop1/Hop1 (mean of both replicates) from that for Hop1-ParB Hop1/Hop1 (mean of both replicates) for the whole genome with Y scale as indicated to illustrate full range of difference and difference in signal across the genome

